# Azurify integrates cancer genomics with machine learning to classify the clinical significance of somatic variants

**DOI:** 10.1101/2025.04.18.649588

**Authors:** Ashkan Bigdeli, Darshan S. Chandrashekar, Akshay Chitturi, Chase Rushton, A. Craig Mackinnon, Jeremy Segal, Shuko Harada, Ahmet Sacan, Robert B. Faryabi

## Abstract

Accurate classification of somatic variations from high-throughput sequencing data has become integral to diagnostics and prognostics across various cancers. However, the classification of these variations remains highly manual, inherently variable, and largely inaccessible outside specialized laboratories. Here, we introduce Azurify - a computational tool that integrates machine learning, public resources recommended by professional societies, and clinically annotated data to classify the pathogenicity of variations in precision cancer medicine. Trained on over 15,000 clinically classified variants from 8,202 patients across 138 cancer phenotypes, Azurify achieves 99.1% classification accuracy for concordant pathogenic variants in data from two external clinical laboratories. Additionally, Azurify reliably performs precise molecular profiling in leukemia cases. Azurify’s unified, scalable, and modular framework can be easily deployed within bioinformatics pipelines and retrained as new data emerges. In addition to supporting clinical workflows, Azurify offers a high-throughput screening solution for research, enabling genomic studies to identify meaningful variant-disease associations with greater efficiency and consistency.

## INTRODUCTION

The classification of clinically relevant genomic variations through next-generation sequencing (NGS) is integral for precision clinical diagnosis and molecular-driven treatment stratification (Morash et al. 2018; Mardis 2019). Professional organizations such as the American College of Medical Genetics (ACMG), American Molecular Pathologists (AMP), American Society of Clinical Oncology (ASCO), and the College of American Pathologists (CAP) have all iterated over systems and guidelines to assist in defining the pathogenicity and reporting criteria for cancer variations (Sukhai et al. 2016; Sirohi et al. 2020). Despite efforts from these groups to establish relevant classification criteria, studies have shown that variability across practitioners persist and only 41% of responding laboratories used published guidelines without any alteration, and 18% of respondents used no published schema (Spence et al. 2019). While data consortiums and professional guidelines aim to assist in variant classification, there is a clear gap in accessible methods that can reduce variability in the field. Additionally, there is no known framework for laboratories to leverage and scale their own classifications to assist evaluation of future events as new data becomes available, which may be particularly useful to those who combine published guidelines or develop their own internal schema.

Today, cancer variant classification relies on the manual review of pertinent resources that help clinical practitioners evaluate a given variant and its association with a particular disease. ACMG and AMP outline their criteria for reporting sequence variants in germline and somatic cancers respectively and additionally provide several resources that can be used to categorize variants based on pathogenicity potential (Richards et al. 2015; Li et al. 2017). While classification schemas may differ in their number of classes, nomenclature, and criteria; these schemas can be summarized broadly as pathogenic variants having clear disease implications, likely pathogenic variants having a high likelihood of disease implication; variants of uncertain significance (VUS) having an unknown impact, and likely benign or benign variants having little or no association with cancer. The criteria to make these classifications include but are not limited to the presence of available therapies, allelic frequency, status in population/germline/somatic variant databases, pathway involvement, predictive software, and available publications and case studies.

To ease the time burden of cross-referencing resources across disparate datasets, variant classification methods and platforms have been developed that aggregate well-defined resources. Platforms such as ClinGen, which aims to aggregate, curate, and disseminate variant curation data have become valuable tools in classifying genomic variants. To extend this further, algorithms have been developed to computationally score and classify variants (AlKurabi, AlGahtani, and Sobahy 2023; Li et al. 2022). Yet, none of the existing methods have leveraged machine learning to directly classify variants by integrating a full feature set from recommended resources with data from a CAP/Clinical Laboratory and Amendments (CLIA) compliant diagnostic center.

To address this, we developed and deployed Azurify, a machine learning-based model trained using over 15,000 variants classified according to a five-tiered schema developed internally at a CAP/CLIA-compliant diagnostic laboratory. This schema categorizes variants into Pathogenic, Likely Pathogenic, VUS, Likely Benign, and Benign, paralleling widely adopted classification systems but based entirely on internal curation standards, workflows, and expert review. Azurify integrates a comprehensive feature set including variant annotations, population frequency data, clinical databases, pathway associations, and predictive scoring algorithms into a unified classification engine. The model is fully pipeline-able, allowing seamless integration into existing bioinformatics workflows. Its architecture is modular, easily retrainable, and designed to scale with expanding datasets, enabling it to evolve alongside newly available clinical or consortium data.

When tested on independent datasets from two external CAP/CLIA-validated laboratories, Azurify achieved a classification accuracy of 99.1% on concordant variants. Additionally, Azurify was able to recapitulate established molecular maps in multiple hematological malignancies. The Azurify algorithm and resources are well documented and modularly designed to allow rapid deployment and adaptation to specific needs. Our analysis shows that the application of Azurify addresses the unmet need for robust and accessible cancer variant classification by providing an effective and accurate variant pathogenicity meta-predictor that could assist as a preliminary screening tool for manual clinical review. Azurify also serves as a high-throughput screening tool for research, particularly in studies seeking to analyze large cohorts and identify meaningful genotype-phenotype associations at scale. By providing standardized and reproducible classifications across thousands of variants, Azurify has the potential to accelerate discovery and hypothesis generation in cancer genomics.

Azurify offers a scalable and accurate variant pathogenicity predictor that addresses key limitations in current variant classification practices. Its modularity, documentation, and availability at https://github.com/faryabiLab/Azurify make it a valuable resource for both clinical and research applications seeking to enhance and scale genomic variant classification.

## Results

### The Azurify Model

Azurify seeks to combine features defined by experts in the field of somatic variant annotation with clinically issued training labels to generate a predictive machine learning classifier (**Figure 1**). We designed Azurify using gradient boosted decision trees (GBDT). This class of algorithms are able to produce accurate results in multi-class classification problems. GBDT algorithms offer high performance when using heterogeneous data types with multiple decision dependencies, which is often the case in cancer variant classification (Prokhorenkova et al. 2018). To train the GBDT model, a 50-50 train-test split was performed on nearly a million variants classified using tumor-only molecular profiling from 8,202 patients at the University of Pennsylvania (UPenn) encompassing 138 electronic health records (EHR) designated cancer phenotypes (**Table S1**). Over 91.16% of variants retrieved from the UPenn cohort were classified as benign, with another 4.17% and 3.78% classified as pathogenic and VUS, respectively. Only 0.03% were classified as likely pathogenic and even fewer, 0.02% were classified as likely benign (**Figure S1A**). Longitudinal analysis of variants that had been classified multiple times showed that 3.12% of variants changed classes, with most variants being re-classified as VUS (**Figure S1B**). These data were fed to the GBDT model at a learning rate of 0.3 and cross-validation testing accuracy showed an average test accuracy of 99.5%, which did not measurably increase after 200 training iterations with the difference between iterations <0.000001% (**Figure S2A**). Iterative model generation using randomly sampled variants from the training set showed that at least 300,000 variants were required to distinguish the sparsely used classes (**Figure S2B**). As a result, the final Azurify model was trained using the breadth of the training set (448,319 variants) and achieved 99.86% training accuracy at 198 iterations (**Figures S2A and S2B**).

**Figure 1.**
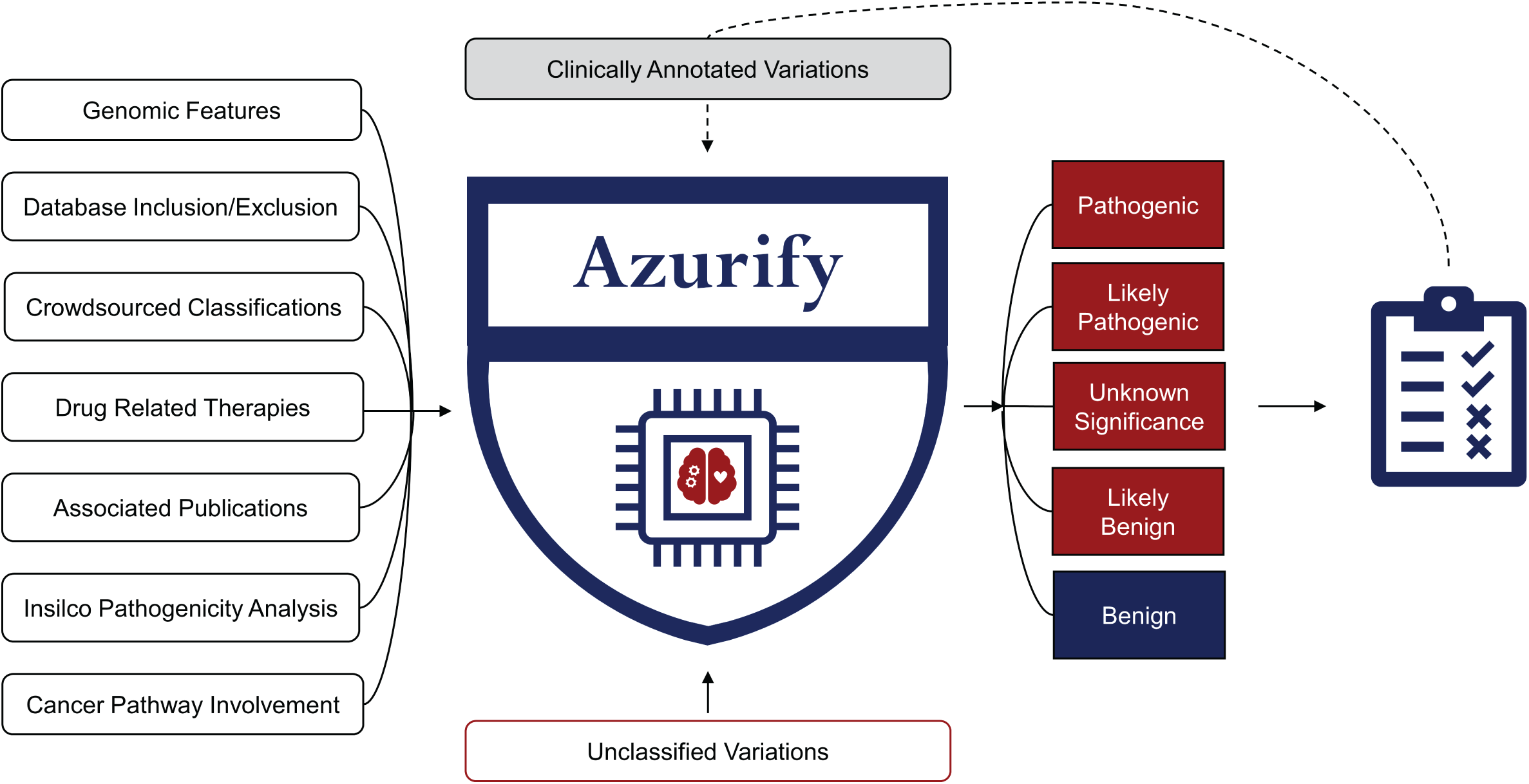
The Azurify Model. Azurify leverages gradient boosted decision trees (GBDT) on a feature set of ACMG/AMP/ASCO/CAP defined resources (left column), which are routinely utilized in the classification of somatic variants. These features are then aggregated and integrated with clinically reviewed variants (top-center gray box) to create a predictive model (center logo) capable of classifying protein changes based on their pathogenicity (right column).

For model features, we have selected 8 resources in conjunction with genomic features to encompass published guidelines as outlined by ACMG and AMP (Richards et al. 2015; Li et al. 2017). This included data for available therapies (Griffith et al. 2017), presence in healthy (Gudmundsson et al. 2022) and disease population (Tate et al. 2019), gene presence in cancer pathways (Kanehisa and Goto 2000), both crowd-sourced (Landrum et al. 2018) and software-derived (Qi et al. 2021) pathogenicity, associated publication data (Allot et al. 2018), and genomic features such as protein domain (UniProt Consortium 2021), translational effect (Cingolani et al. 2012), exon, variant type, and allelic frequency (**Table S2**). To assess the impact of each resource, feature importance was calculated as the average percentage of contribution to each of the 5 classes in our multi-classification model. This analysis revealed that each of the resources recommended through professional guidelines effectively contribute to variant classification (**Figure 2A**). One-vs-Rest receiver operator curves (ROC) showed that Azurify outperformed any single resource for classification of pathogenic and VUS variants (**Figures 2B and 2C**). Similarly, Azurify outperformed any single resource for classification of likely pathogenetic, likely benign, and benign variants (**Figure S3**).

**Figure 2.**
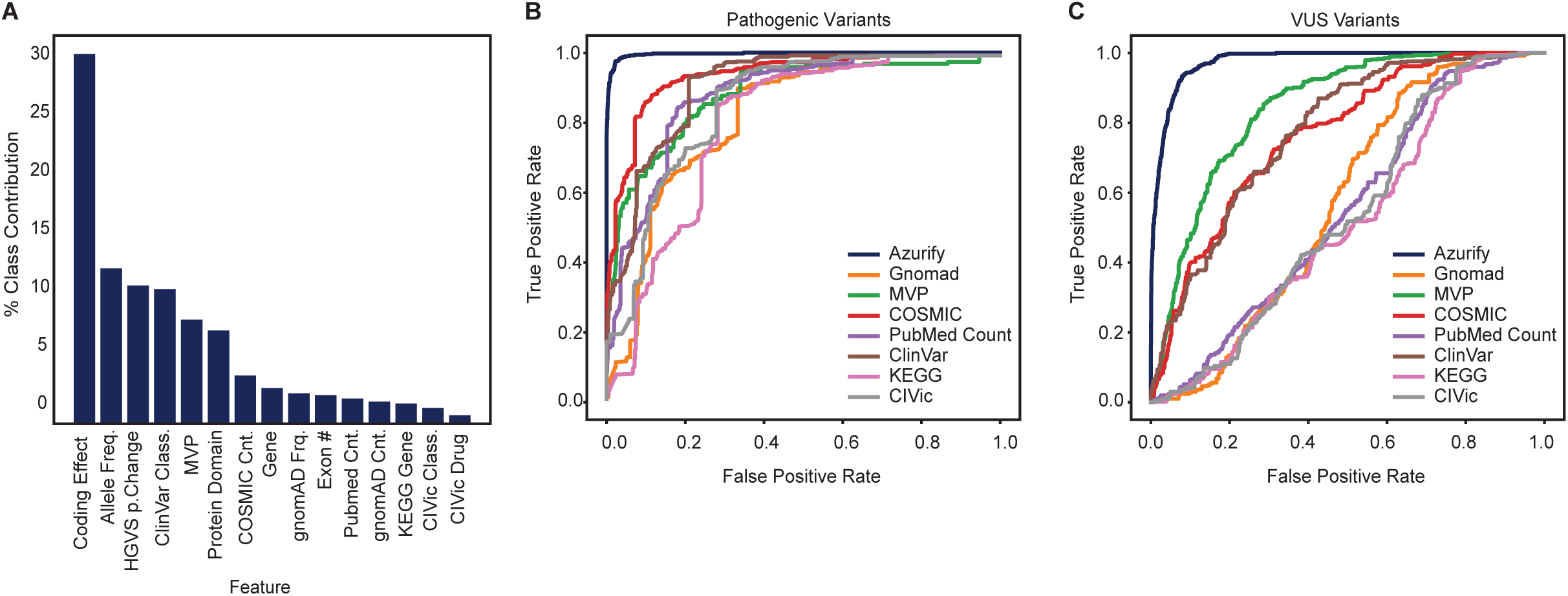
Azurify’s individual feature importance and performance. **A**: Calculated feature importance shows translational effects, crowd sourced classifications, and *in silico* predictors are among the most influential features. Barplot showing percentage class contribution as a function of annotation features. Note that drug targets and pathway involvement contribute below 5% to model classification, which may be explained by the relatively low number of available therapies, as well as targeted assay bias towards cancer genes. **B, C**: ROC curves of each feature’s ability to classify variants in the holdout data. Each ROC curve shows performance evaluation in predicting pathogenic (B) and VUS (C) variants when a given knowledgebase (i.e, gnomAD, MVP, etc) is added to genomic features (i.e, domain, coding effect, etc). Azurify outperforms any individual knowledgebase.

### Azurify accurately classifies variants in the holdout data

To examine Azurify performance, we first compared concordance between Azurify and manually annotated variants in holdout data with the same genes and loci used in model training (**Figures 3A and 3B**). Benign variations encompass over 91% of the variations found in the holdout dataset. While these variations are not clinically relevant in cancer diagnostics, their prevalence requires accurate segregation. Azurify effectively achieved this objective by accurately classifying 99.89% of benign variations (**Figure 3A**). VUS, where the clinical impact cannot be fully ascertained, represented the next biggest variant class in the holdout dataset. Azurify achieved 95.63% accuracy in VUS variant classification. Likely benign and likely pathogenic variants were classes that were infrequently observed in our available data, representing less than 0.06% of clinical classifications. Given the paucity of these classes in both holdout and training datasets, Azurify only showed 47.74% and 52.76% accuracy in classification of likely benign and likely pathogenic variants, respectively. Pathogenic variants are the most impactful as they are likely to affect prognosis and treatment, having an established role in cancer. Notably, Azurify correctly classified pathogenic mutations with 98.81% accuracy when compared to manual clinical review (**Figure 3A**). The achieved accuracy in the holdout set indicates that Azurify performs well in pathogenic, VUS, and benign categories, which are used frequently in clinical practice, but encounters potential challenges when applied to likely pathogenic and likely benign classifications, which are infrequently applied in clinical practice.

**Figure 3.**
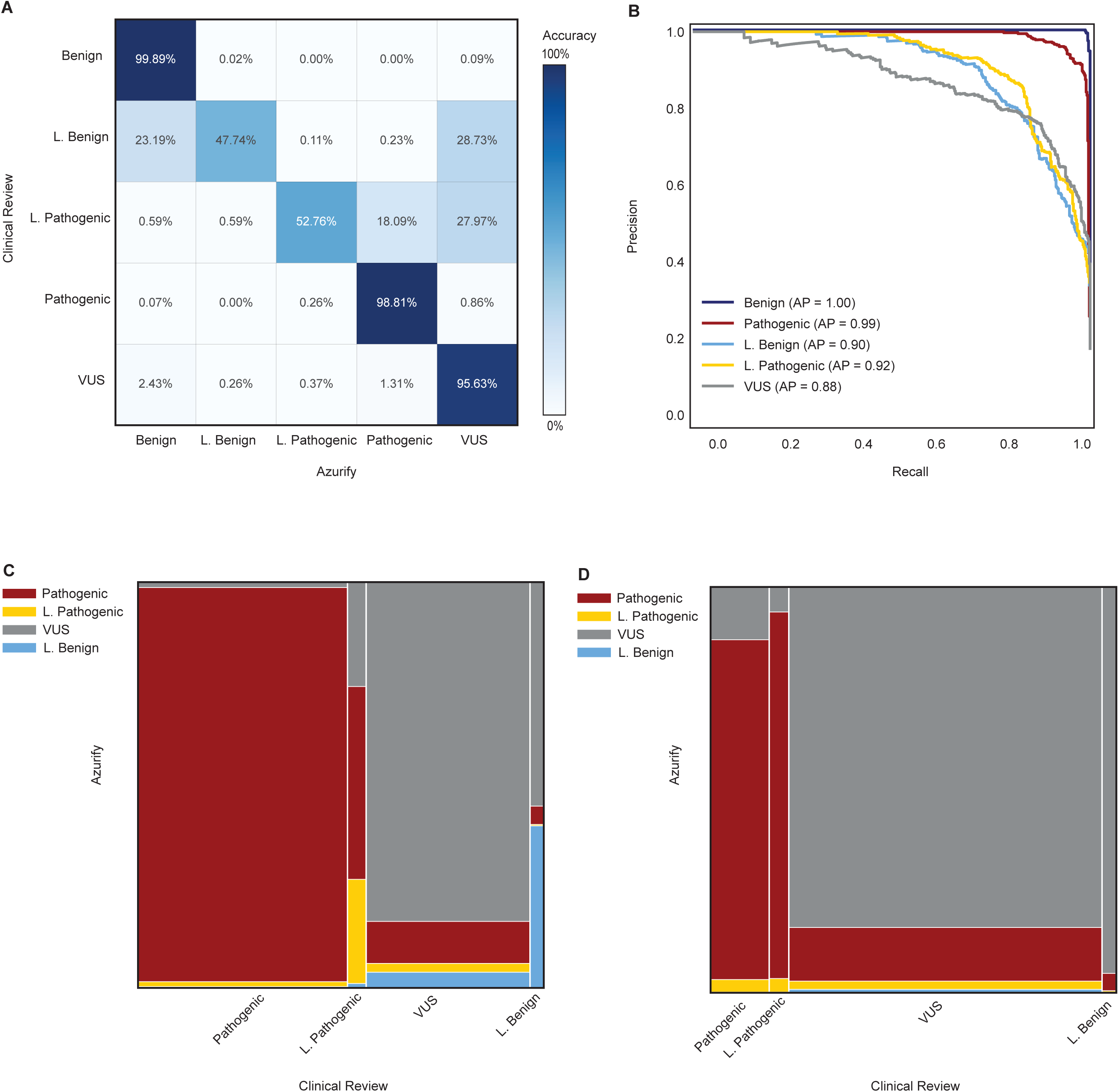
Evaluation of Azurify performance with holdout and the UPenn independent datasets. **A**: Azurify accurately classified pathogenic, VUS, and benign variations. Heatmap showing percentage of concordance between Azurify prediction and clinically reported somatic variants for benign, likely benign (L. Benign), likely pathogenic (L. Pathogenic), pathogenic, and VUS variations. Low concordance is observed in likely benign and likely pathogenic categories which had limited training data. **B**: Average precision (AP) shows high accuracy of Azurify in classifying pathogenic and benign variants with VUS variant classification at 88% AP. Each curve shows AP for benign, likely benign (L. Benign), likely pathogenic (L. Pathogenic), pathogenic, and VUS variations. Lower precision is observed in less prevalent classes (likely benign, likely pathogenic). **C, D**: Mosaic plots, in which the width of the bar represents class proportion, shows high accuracy in predicting frequently reported variant classes in the UPenn independent dataset when evaluating the genes present (C) and not present (D) in model training.

Average precision (AP) calculations across the same holdout data showed that Azurify classified pathogenic and benign variants with greater than 99% precision (**Figure 3B**). Classification of VUS variants resulted in an AP of greater than 88%, while classification of pathogenic and likely benign events resulted in 92% and 90% AP, respectively (**Figure 3B**). To examine the effect of cancer type prevalence on performance, we divided the holdout data into quartiles of solid tumor types based on presentation frequency and assessed the average precision (**Table S3**). This analysis showed that the classification of well represented pathogenic, VUS, and benign classes was invariant to cancer type prevalence in the cohort, while the performance varied for less represented likely pathogenic and likely benign classes (**Figures S4A-D**). For example, we did not observe a marked difference between Azurify’s average precision for classifying variants in genetically well characterized and moderately represented breast cancer compared to other solid tumors in the UPenn holdout data (**Figures S4E and S4F**).

### Azurify accurately classifies pathogenic variants independent of clinical NGS assay

To further evaluate Azurify performance, we created an independent test set by obtaining reportable variation data from 3,411 patients sequenced using an independent CAP/CLIA-validated clinical assay at UPenn. This test data was generated using different biochemistry, technology, informatics, and gene sets than those used in the training (**Figure 2**) and validation (**Figures 3A and 3B**), illustrating independence from upstream variables common in clinical cancer diagnostic assays. In addition to the 237 genes included in Azurify training, the UPenn independent dataset comprised 32 new genes that had not been previously observed by the Azurify model (**Table S4**). We evaluated the accuracy of Azurify when compared to clinical review using the same classification criteria.

In the 237 genes for which Azurify had training data, pathogenic variants were classified with an accuracy of 96.52% (**Figure 3C**). When evaluating classes comprising the remaining reportable variants, the accuracies were as follows: 40.18% for likely benign (n = 331), 84.27% for VUS (n = 4183), and 55.32% for likely Pathogenic (n = 430) (**Figure 3C**). Just as in model training, likely pathogenic and likely benign classifications were present infrequently, comprising only 3.1% and 7.1% of variants, respectively. Evaluating the 32 genes for which Azurify had no training data, class accuracy values were as follows: pathogenic 84.26 % (n = 108), likely pathogenic 3.03% (n = 33), VUS 84.59% (n = 585), and likely benign 96% (n = 25) (**Figure 3D**). Publicly available data for the 32 genes evaluated with no prior training data was considerably sparser, explaining the drop accuracy for a model that aggregates such resources. Overall, these analyses show that Azurify performs best when evaluating genes used in training, while still providing acceptable performance in previously unobserved genes.

### Azurify accurately classifies pathogenic variants in datasets from two external laboratories

To ensure the generalizability of Azurify performance, we next examined its ability to classify variants from laboratories outside UPenn. To this end, we obtained variant classification data for patients with Acute Myeloid Leukemia (AML) and lung cancer from clinical laboratories at the University of Alabama at Birmingham (UAB) and the University of Chicago (UC). Similar to UPenn, CAP/CLIA-certified clinical laboratories at UAB and UC perform high throughput sequencing for cancer diagnostics. Each clinical laboratory had independently developed sequencing assays, informatics, and variant classification schemas.

In total, we analyzed 101 clinical cases from UAB and UC (**Table S5**). UAB uses a three-class variant reporting schema, classifying variants as pathogenic, VUS, and benign. In contrast, UC classifies variants into four tiers: pathogenic (Tier 1), likely pathogenic (Tier 2), VUS (Tier 3), and benign (Tier 4). We first evaluated Azurify performance in annotating UAB and UC datasets separately without adjusting for differences between their variant reporting schemas. Across both datasets, Azurify achieved an average classification accuracy of 96.34% and 87.31% for pathogenic variants in lung cancer and AML cases, respectively (**Figures S5A-D**).

Next, we assessed Azurify performance in comparison to clinical reviews while mitigating differences in the UAB and UC variant reporting schemas. We grouped the variants that were originally reported as pathogenic or likely pathogenic into a single pathogenic class. Similarly, we grouped the variants that were originally reported as VUS or benign as the non-pathogenic class. Using these harmonized variant labels from the reporting laboratories as ground truth, we then assessed Azurify performance in classifying pathogenic and non-pathogenic variants from UAB and UC. Azurify achieved a classification accuracy of 98.93% and 92.31% for non-pathogenic and pathogenic variants, respectively, in lung cancer cases from both laboratories (**Figure 4A**). Similar analysis of AML variants from both laboratories showed that Azurify classified non-pathogenic and pathogenic variants with 99.03% and 88.34% accuracy, respectively (**Figure 4B**).

**Figure 4.**
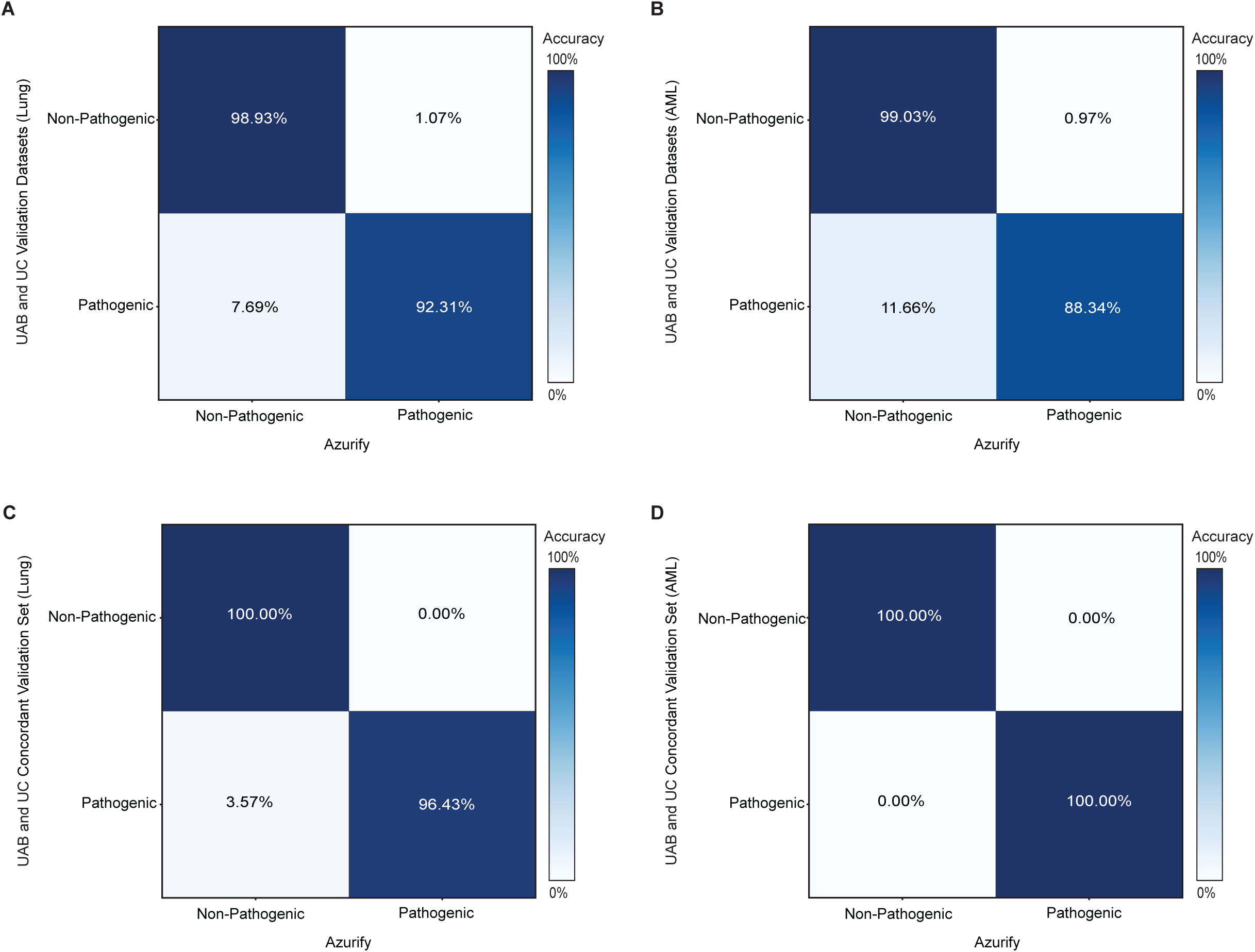
Azurify performance evaluation with independent harmonized data from two external laboratories. **A**: Heatmap showing the Azurify classification accuracy for clinically reported somatic variants (n=5,615) in 50 lung cancer cases from UAB and UC after label harmonization. **B:** Heatmap showing the Azurify classification accuracy for clinically reported somatic variants (n=9,110) in 51 AML cases from UAB and UC after label harmonization. **C**: Heatmap showing the Azurify classification accuracy for lung cancer variants concordantly reported by UAB and UC (n = 31). **D**: Heatmap showing the Azurify classification accuracy for AML variants concordantly reported by UAB and UC (n = 954).

Close examination of UAB and UC datasets revealed a sparsity of overlap between their reported variants, potentially due to differences in variant reporting schema (See Methods). Nevertheless, we assessed the level of concordance between UAB and UC (**Figures S5E and S5F**) and examined Azurify performance based on concordantly reported variants (**Figures 4C and 4D**). When evaluating concordantly reported variants, Azurify achieved 100% accuracy in classifying non-pathogenic events in both AML and lung cancer. Similarly, Azurify exhibited high accuracy in identifying concordantly reported pathogenic variants with a classification accuracy of 96.4% and 100% in lung cancer and AML cases, respectively.

Taken together, these benchmarking analyses support Azurify model generalizability and show its ability to robustly classify pathogenic variants despite reporting schema differences. These studies further suggest that data harmonization could enhance Azurify performance and hints to classification schema standardization as a key step towards developing more reliable variant classification tools.

### Azurify compares favorably against another variant classification tool

CancerVar is a tool for the clinical interpretation of somatic variants (Li et al. 2022). While Azurify does not attempt to interpret somatic variations, both methods do classify variants according to professional guidelines and thus warrant comparison.

We first compared performance of CancerVar and Azurify using the UPenn independent dataset. CancerVar uses AMP/ASCO/CAP 2017 reporting guidelines, which classify variants into 4 tiers; hence, there is no exact 1:1 conversion between CancerVar and Azurify classifications. Nonetheless, the UPenn independent dataset (**Figures 3C and 3D)** contained only the first 4 variant classifications allowing for the following conversion: Tier 1 = Pathogenic, Tier 2 = Likely Pathogenic, Tier 3 = VUS, and Tier 4 = Likely Benign variants. Using CancerVar API, we attempted to annotate the entirety of variants in the UPenn independent dataset based on their hg19 assembly chromosome, genomic position, reference, and alternate allele. However, the CancerVar API failed to produce results for 3,004 out of 11,080 queried variants (27.1%), forcing us to remove them from the comparative analysis.

Azurify outperformed CancerVar in detection of pathogenic, VUS, and likely benign variants but not likely pathogenic (**Figure 5A**). CancerVar showed higher accuracy in annotating likely pathogenic variants with an accuracy of 41.03% compared to Azurify’s 24.10%, potentially due to the low number of this class of variants in Azurify training dataset. Notably, Azurify’s classification of pathogenic variants was 7.1 times more accurate than CancerVar (97.51 % vs 13.6%). Similarly, Azurify produced 2.1-fold (38.8% vs 18.3%) and 1.2-fold (84.31% vs 67.11%) higher accuracy in classification of likely benign and VUS mutations, respectively.

**Figure 5.**
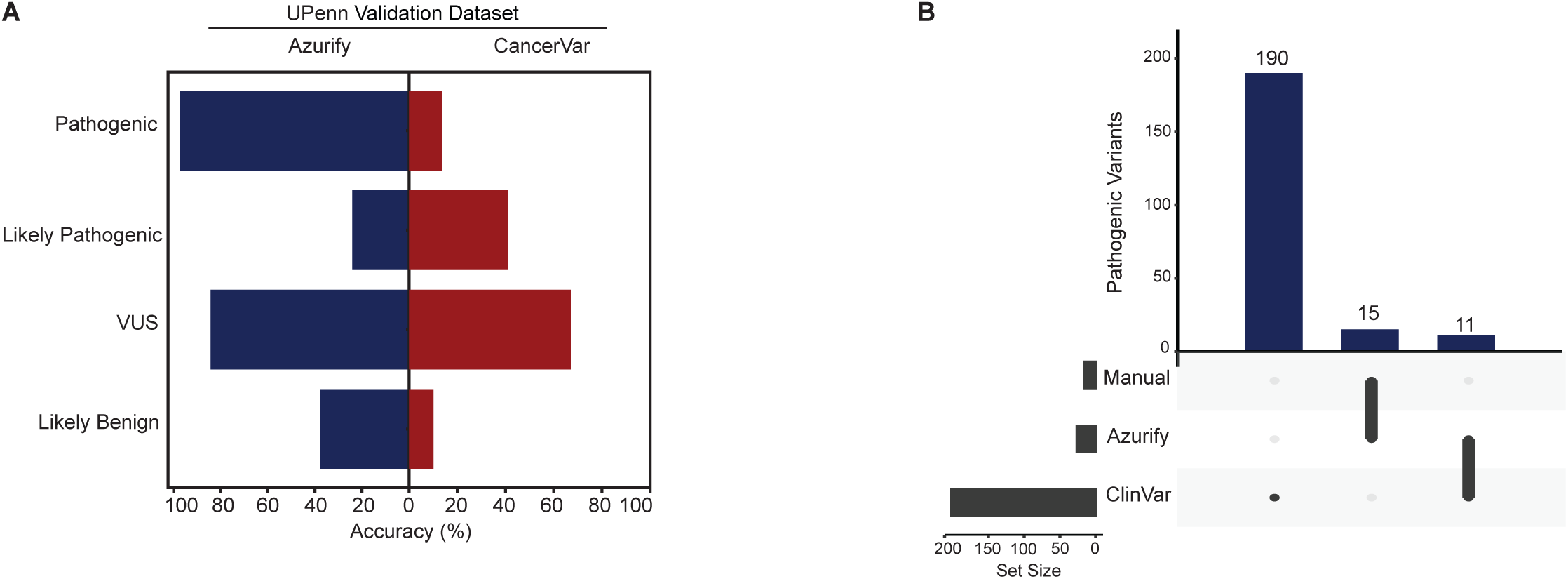
Azurify outperforms CancerVar. **A**: Azurify compares favorably to CancerVar. Azurify (blue) performs well when compared to CancerVar (red) when using the UPenn independent test data to evaluate % classification accuracy (X-axis). Notably pathogenic variants classified at a significantly higher rate in Azurify. Additionally, all variants return a classification when using Azurify where CancerVar fails to classify 27% of input data. **B**: Azurify identification of emergent variants. Up-set analysis shows the overlap of variants from the UPenn independent test set clinically reviewed as likely pathogenic but classified as pathogenic by Azurify and ClinVar. As discussed in the main text, variants classified by both Azurify and ClinVar as pathogenic were further reviewed and contained a somatic cancer variant required for clinical trial inclusion as well as pathogenic germline variant not associated with cancer.

We next compared performance of CancerVar and Azurify using all the variants in the UAB and UC datasets (**Figures 4A and 4B**). This analysis showed that Azurify maintained its performance advantage. While Azurify correctly classified 92.2% of all the pathogenic variants in the two datasets, CancerVar completely failed to process 30.02% of the variants and only correctly classified 52.54% of the variants that were processed (**Figure S6A**). Finally, we benchmarked relative performance of CancerVar and Azurify using variants concordantly reported by both UAB and UC (**Figures 4C and 4D**). This analysis led to markedly closer performance, with Azurify and CancerVar classifying pathogenic variants with an accuracy of 98.53% and 95.56%, respectively (**Figure S6B**). Taken together, this data demonstrates improved performance of Azurify compared to CancerVar and further suggests the benefit of variant schema harmonization in improving variant classification tools.

### Azurify identifies emergent somatic variants and established germline variants

When reviewing Azurify-classified data, we observed several variants that were algorithmically labeled as pathogenic while clinically reported as likely pathogenic (**Figures 3C and 3D**). To corroborate this assertion, we evaluated 190 ClinVar classified pathogenic variants and assumed the crowdsourced classifications as ground truth. Of these 190, 15 were classified as pathogenic by both Azurify and manual review and 11 were classified as pathogenic only by Azurify and ClinVar (**Figure 5B**). Within these 11 overlapping variants, Azurify classified a p.Val834Leu variation in EGFR as pathogenic. At the time of assessment, this variation was an inclusion criteria in a phase 1 clinical trial, underscoring Azurify’s ability to identify relevant somatic variations based on emerging literature (Clinical trial: NCT04085315).

Azurify was also able to identify causal germline variants not associated with cancer according to ClinVar. Variant p.Arg790Gln in gene SMC1A was annotated by two institutions, 7 years apart, as being of both germline origin and pathogenic (NCBI ClinVar query). This variant is relevant in congenital muscular hypertrophy-cerebral syndrome (Deardorff et al. 2007) and explains its downgraded clinical classification when reported by a somatic diagnostic center.

### Azurify accurately recapitulated patterns of co-mutation in Acute Myeloid Leukemia

To showcase Azurify’s effectiveness in identifying prognostically relevant AML subtypes, we analyzed genomic variants across 326 AML patients, spanning 617 sequencing events in the UPenn cohort. AML is a hematologic malignancy with prognostically important molecular subtypes (DiNardo and Cortes 2016). Specific subtypes can be determined based on the presence or absence of cooperative variations within epigenetic regulators and DNA-binding transcription factors.

In line with earlier reports (Park et al. 2020), Azurify identified *DMNT3A* pathogenic variations in 23.7% of the cases in the UPenn AML cohort (**Figure 6A**). Azurify also correctly detected the expected frequency of co-mutations in *DMNT3A* and *FLT3* as well as *NPM1* and *FLT3* (Bezerra et al. 2020) (**Figure 6A**). Further concordance between published literature and Azurify pathogenicity classification was observed when we examined mutations in RAS proto-oncogenes within the UPenn AML cohort. RAS variations have been reported in 10-15% of AML patients (Al-Kali et al. 2013). Azurify mirrored the reported mutation rate by classifying 11.5% of UPenn AML patients as RAS mutated. More specifically, NRAS (G12 C/D/S, G13 C/D/R/V, T581, Q61) and KRAS (G12 A/D/S/V, G13D, Q61H, R68S, K117N) mutations were reported in 11.6% and 4.7% of AML cases, respectively (Ball et al. 2021). Our analysis of the UPenn AML cohort with Azurify recapitulated these observations and identified 12.72% and 6.5% mutation rates in these residues of NRAS and KRAS, respectively (**Figure 6A**).

**Figure 6.**
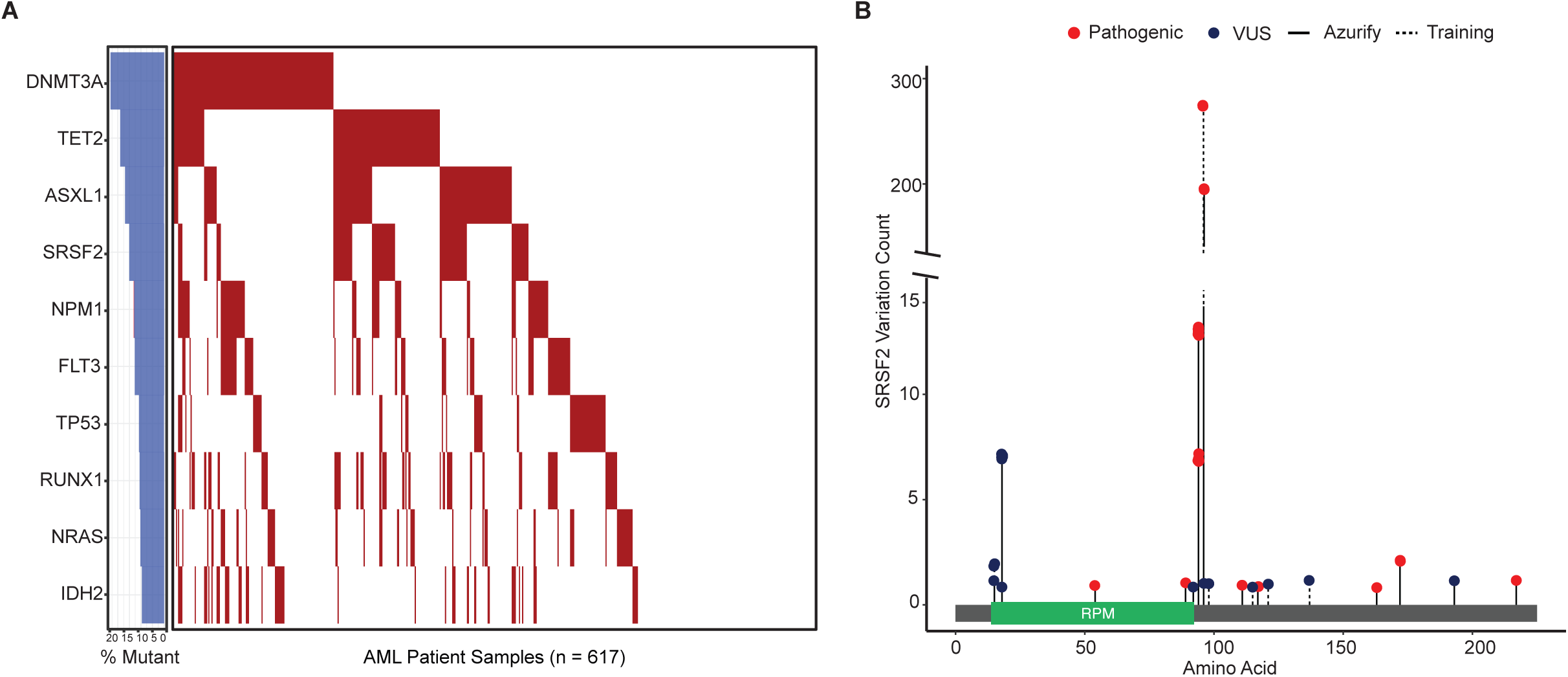
Azurify accurately profiles co-occurring mutations in the UPenn AML cohort. **A**: Co-mutation analysis of the UPenn AML cohort using Azurify. Barplot on the left shows mutation frequencies for noted genes in the UPenn AML cohort. Heatmap on the right shows pathogenic mutations per patient (x-axis) as predicted by Azurify. **B**: Lollipop plot of SRSF2 mutations classified as VUS and pathogenic by Azurify. Plotted SRSF2 variation counts (y-axis) show that training data derived from targeted sequencing is centered at a particular amino acid (x-axis), 95, a common variant in AML patients. Despite lacking training data from sequences flanking the SRSF2 hotspot mutation, Azurify is still able to classify pathogenic and VUS variants.

This analysis also confirmed our studies in Figure 5B and showed that Azurify can accurately identify mutations outside of the training regions of the genes for which it was trained. Despite training data being centered around variations at amino acid 95 of SRSF2, a common occurrence in AML patients (Grimm et al. 2021), Azurify successfully classified oncogenic variations in SRSF2 by identifying variations in flanking nucleotides coding for the amino acid 95 (**Figure S7A**), enabling accurate evaluations beyond its training loci in known cancer genes (**Figure 6B**).

### Azurify accurately identifies mutations in Chronic Lymphocytic Leukemia

To further evaluate Azurify’s classification ability, we examined mutations in 73 chronic lymphocytic leukemia (CLL) patients in the UPenn cohort, which in comparison to AML had a much lower prevalence in our cohort (**Table S6).** To this end, we compared the mutation frequency of 22 genes mutually examined in both UPenn and Kinsbacher *et al*. cohorts (Knisbacher et al. 2022). Notably, we observed no significant differences in the frequency of pathogenic mutations identified by Azurify and Kinsbacher *et al*. (**Figure 7A**, Welch’s t-test P-value=0.227).

**Figure 7.**
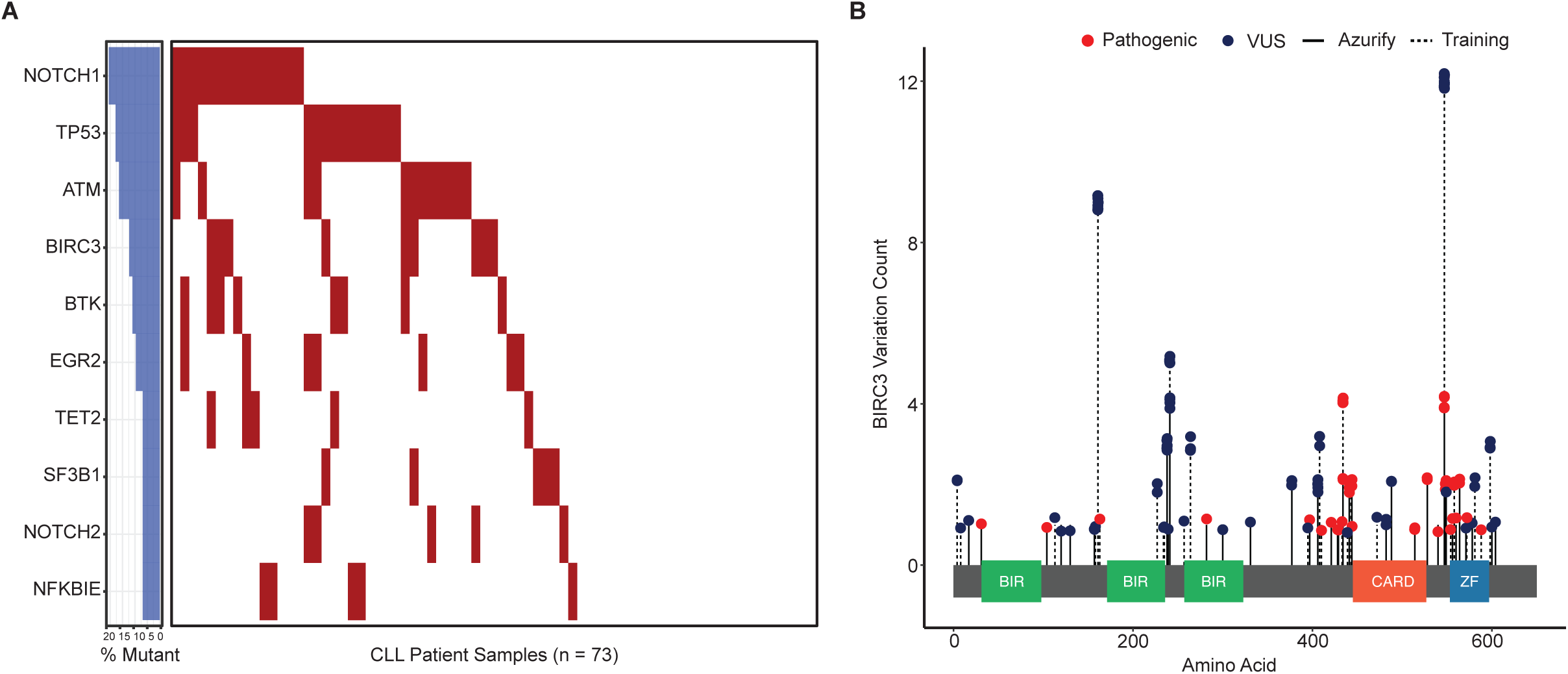
Azurify recapitulates frequency of pathogenic mutations using the UPenn CLL cohort. **A**: Mutation analysis of the UPenn CLL cohort using Azurify. Barplot on the left shows mutation frequencies for noted genes in the UPenn CLL cohort. Heatmap on the right shows pathogenic mutations per patient (x-axis) as predicted by Azurify. **B**: Lollipop plot of BIRC3 mutations classified as VUS and pathogenic by Azurify. Plotted BIRC3 variation counts (y-axis) show that training data derived from targeted sequencing is centered around ZF domain variant in CLL patients. Despite lacking training data from sequences flanking the ZF domain, Azurify is still able to accurately classify pathogenic and VUS variants.

Upon closer examination of *BIRC3*, a gene associated with unfavorable outcomes in CLL (Diop et al. 2020; Tausch and Stilgenbauer 2020), we again observed Azurify’s ability to accurately classify pathogenic variants in amino acids of known cancer genes, which are reported to be pathogenic (Diop et al. 2020) but were not present in the Azurify’s training data (**Figures 7 and S7B**).

## Discussion

Azurify benchmarking analysis revealed the promise of leveraging resources recommended in professional guidelines in conjunction with expert annotations to build classification models for efficient and accurate labeling of pathogenic variants in cancer. Our studies showed that Azurify can reach up to 96% accuracy when evaluating pathogenic variants in datasets from three independent CAP/CLIA-certified laboratories. Notably, Azurify reached 99.1% overall accuracy in classifying variants concordantly reported by two institutions. Additionally, we showed Azurify’s ability to effectively recapitulate molecular profiles observed in AML and CLL patients. Lastly, our method shows potential in profiling cancers for emergent variants including those informing clinical trials.

With Azurify showing promise, the discussion of extending the classification data used in model training beyond a single institution is required. The SOCIAL project conducted in 2019 evaluated the application of published guidelines and found that 59% of respondents did not adhere to published schema without alteration and concluded that aligning classification methods would reduce variation across reporting laboratories (Spence et al. 2019). Future iterations of Azurify hope to fill this unmet need by training models using a variety of classification methods across a broader range of clinical laboratories. The problem of circular labeling, where false positives are incorrectly confirmed and persist through data and derived models, is a consideration in multi-institutional training when using supervised machine learning. The issue of false classification through circular labeling is considered by Cheng *et al*. as they approach pathogenicity classification through predictive protein structures (Cheng et al. 2023). While impact based on predicted structure is not a professionally recommended feature, its emergence and accuracy may boost model performance and mitigate inherent circular labeling when using publicly available data for model generation. Other features such as clinical trial data and pharmacogenomic data may also boost model performance and warrant future work.

To maximize algorithmic identification of emergent cancer-associated variants, we can benefit from Azurify’s flexibility, which allows continuous model improvement and publishing. For instance, MLops, which operationalizes constant integration and deployment of machine learning models, can be applied to Azurify so that new somatic variant classifications as well the latest resource data (i.e, new gnomAD or COSMIC releases) can be used to iteratively publish models with higher performance to more accurately reflect the latest observations in the field.

In conclusion, Azurify achieves high accuracy and attempts to address the known gap in accessibility, variability and reproducibility of variant classifications through the application of machine learning using clinically classified variants. Its modular design allows users to classify variants with the existing model or newly trained models. Azurify is well documented and can be easily installed as a standalone program through https://github.com/faryabiLab/Azurify.

## Acknowledgments

We would like to express our sincere gratitude to the Center for Personalized Diagnostics for their invaluable support and resources, which significantly contributed to the success of this research. Special thanks to Dr. Uri Hershberg for his guidance in the initial steps of the Azurify project. This work was supported in part by R01-CA230800, R01-CA248041, and U01DK123716 (to R.B.F.), and Human Pancreas Analysis Program through U01-DK112217, U01DK123594.

## Authors Contributions

Conceptualization: A.B., R.B.F.; Methodology: A.B., R.B.F.; Investigation: A.B., R.B.F.; Formal Analysis: A.B., A.S., D.S.C., A.C., C.R., R.B.F., Resources and Reagents: A.C.M., J.S., S.H.; Writing-Review & Editing: A.B., A.S., R.B.F.; Writing-Original Draft: A.B., R.B.F.; Funding Acquisition: R.B.F.; Supervision: R.B.F.

## Code Availability

All the code needed to evaluate the conclusions in the paper is publicly available at https://github.com/faryabiLab/Azurify.

## MATERIALS AND METHODS

### UPenn Dataset

To train the Azurify variant classification algorithm, we extracted tumor-only high throughput sequencing data from 8,202 patient samples from the Hospital of the University of Pennsylvania’s (HUP) Center for Personalized Diagnostics (CPD) genomic database. Data were extracted and selected for single nucleotide variations (SNVs) and small insertions and deletions (indels) that were classified and reported according to a variation of ACMG/AMP 2015 guidelines. This yielded 896,899 variations sequenced over a 6-year period with 25,789 variations being unique and dispersed across 248 cancer consensus genes. These data were then queried against the HUP electronic medical record system and found to encompass 138 distinct cancer phenotypes according to input histology. The resulting dataset was then de-identified through selection of features relevant for classification.

### UAB and UC Datasets

To evaluate the performance characteristics of the Azurify variant classification algorithm, we also obtained data from two laboratories that perform tumor-only high throughput sequencing for precision cancer medicine at the University of Alabama at Birmingham (UAB) and the University of Chicago (UC). Each institution provided de-identified variant data annotated with pathogenicity classifications by their respective team of experts. The resulting dataset contains 14,725 variants annotated with classification labels generated by both institutions and Azurify.

UAB provided variant data from 31 AML patients and 30 lung cancer patients. The UAB AML dataset contained a total of 2,650 variants that spanned 53 distinct genes. UAB experts classified these AML variants as being either pathogenic, VUS, or benign. The UAB lung cancer dataset contained a total of 361 variants that spanned 211 genes. Only pathogenic and VUS variants were provided by UAB for the lung cancer dataset.

UC provided variant data from 20 AML patients and 20 lung cancer patients. The UC AML dataset contained a total of 6,460 variants that spanned 150 distinct genes. The UC lung cancer dataset contained a total of 5,573 variants that spanned 158 distinct genes. UC experts classified AML and lung cancer variants into four tiers: pathogenic (Tier 1), likely pathogenic (Tier 2), VUS (Tier 3), and benign (Tier 4).

To harmonize and compare UC and UAB cohorts, classifications were grouped as follows: pathogenic (pathogenic and likely pathogenic) and non-pathogenic (VUS and benign). Of the 53 UAB AML genes and 150 UC AML genes, 38 were found to be shared. Of the 38 shared UAB/UC AML genes, 24 genes in 52 patients contained alterations that had matching clinical classifications. Within these 24 genes and 52 patients there were 954 variants that shared a clinical classification between the two institutions, with 48 of these variants being unique. Of the 211 UAB lung cancer genes and 159 UC lung cancer genes, 75 were shared. Of the 75 shared UAB/UC lung cancer genes, 6 genes in 24 patients contained alterations that had matching clinical classifications. Within these 6 genes in 24 patients, there were 31 variants that shared a clinical classification between the two institutions, with 9 variants being unique.

### Feature Engineering

We have selected 8 resources in conjunction with genomic features (i.e, translation effect, allelic frequency) to encompass published guidelines for the classification of somatic variations in cancer (**Table S2**). Available therapies for specific alterations were extracted from the Clinical Interpretation of Variants in Cancer database, CIViC, via the accepted variants data release variant call file (vcf). Genomic features such as allele frequency, amino acid change, variant type, exon number, and effect were acquired through vcf annotation with SnpEff version 4.1.1 (Cingolani et al. 2012). To determine population prevalence, allelic counts and frequencies were extracted from Genome Aggregation Database, gnomAD, accessible vcf version 2.1.1 (Gudmundsson et al. 2022). Similarly, allelic counts for a given alteration were obtained from the Catalog of Somatic Mutations, COSMIC, via the downloadable vcf v67 (Tate et al. 2019). Missense Variant Pathogenicity prediction software (MVP) was used as an *in-silico* method due to its demonstrated higher AUC scores compared to other predictive models (Qi et al. 2021). To further assist in determination of pathogenicity, clinical assertions were obtained from ClinVar’s variant summary data as of June 2021 (Landrum et al. 2018). Pathway involvement was derived from genes in the KEGG cancer pathway (Kanehisa and Goto 2000). Domain data was also provided to the algorithm through the Uniprot consortium (UniProt Consortium 2021). Lastly, the number of publications associated with a given protein change was derived from National Center for Biotechnology Information’s (NCBI) LitVar application, which allows for the fuzzy searching of publications associated with protein changes (Allot et al. 2018). Data was then aggregated through queries of chromosome, position, reference, and alternate alleles in human genome version 19 (hg19) and effectively forms a comprehensive feature set of resources grounded in professional recommendations. The 8 resources combined with genomic features comprises a set of 15 mixed data types.

### Model Training

We used the Catboost GBDT library (Prokhorenkova et al. 2018). To train the GBDT model, a 50-50 train-test split was performed and fed to the model at a learning rate of 0.3. Cross-validation testing accuracy showed an average test accuracy of 99.5%, which did not measurably increase after 200 training iterations with the difference between iterations < 0.000001%. Iterative model generation using randomly sampled variants from the training set showed that at least 300,000 variants were required to distinguish the sparsely used classes. As a result, the final Azurify model was trained using the breadth of the training set (448,319 variants) and achieved 99.86% training accuracy at 198 iterations.

To rigorously assess model generalizability, key features such as gene, amino acid change, and specific amino acid properties were intentionally withheld during training and evaluated independently. While their exclusion resulted in only minor decreases in overall accuracy, their subsequent inclusion in the final model ensured optimal performance by leveraging features that are reliably available at inference time.

### Analyses and Visualization

Statistical differences between CLL molecular profiles were determined through Welch’s t-test using R (version 4.2.3) (R Core Team 2021). Co-occurrence and gene frequency analysis in AML and CLL were generated using GenVisR (Skidmore et al. 2016). Remaining figures were generated using ggplot2 and extended tidyverse packages (Wickham 2016).

### Software Usage

Azurify is a Python based command line tool that requires the chromosome, position, reference, alternate base, as well as the allelic frequency in a tabular format as input. Azurify runs this input using SnpEFF version 4.1.1 to obtain annotations using hg19 as a base genome build. If required, the user may also select the hg38 genome build and a conversion tool, liftover 1.1.16, will be used to convert annotations and any failed conversions will be displayed in the user’s designated output folder directory. After user input into Azurify, resource aggregation is performed using the pandas python library. The completed dataset is then fed into Azurify model to process and obtain classifications, which are then directly merged back to the input creating the final call set. The user will receive all model features, variant classifications and the prediction probability of each variant class as tabular output. Software and its accompanied documentation can be found at https://github.com/faryabiLab/Azurify.

## Supplementary Information

**Figure S1.**
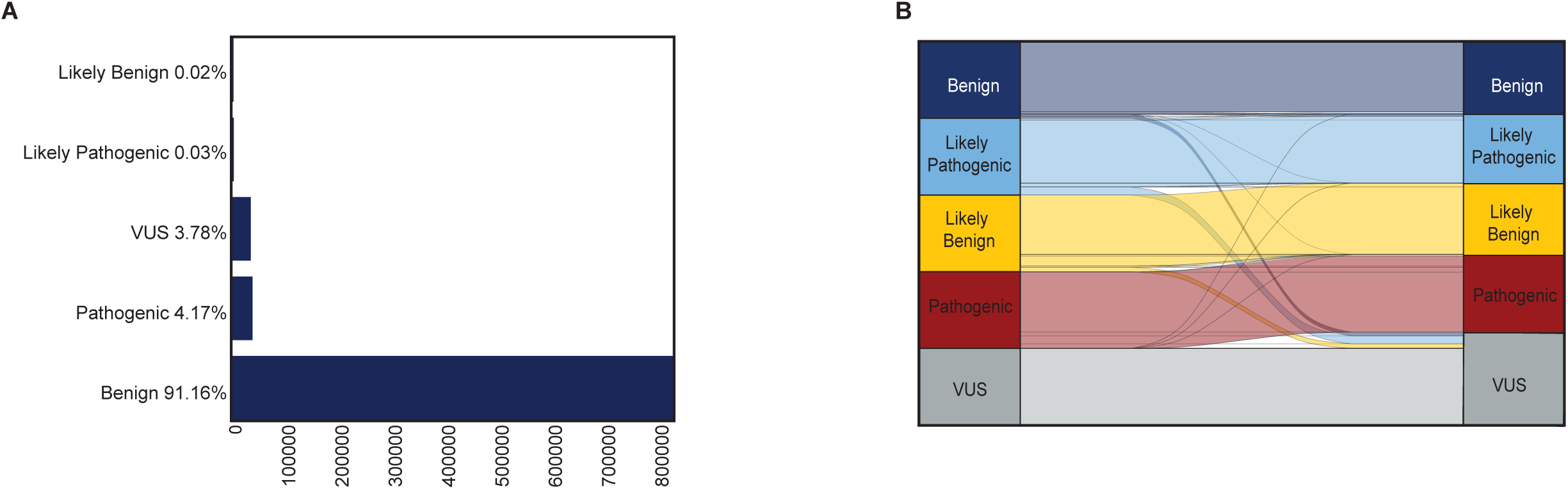
Characterization of different variant classes in the Azurify training data. **A**: Barplot shows class distribution of variant classes. A majority of variants in the training data are classified as benign with pathogenic and VUS classifications observed between 3.7% and 4.2% of the time, respectively. Likely pathogenic and likely benign classifications are infrequent in the training data and only observed in less than 0.05% of variants. **B**: Alluvial analysis, which shows the flow of reclassified variants in the clinic, indicates that variant reclassification occasionally occurred during manual annotation in the training data, mostly impacting VUS variants.

**Figure S2.**
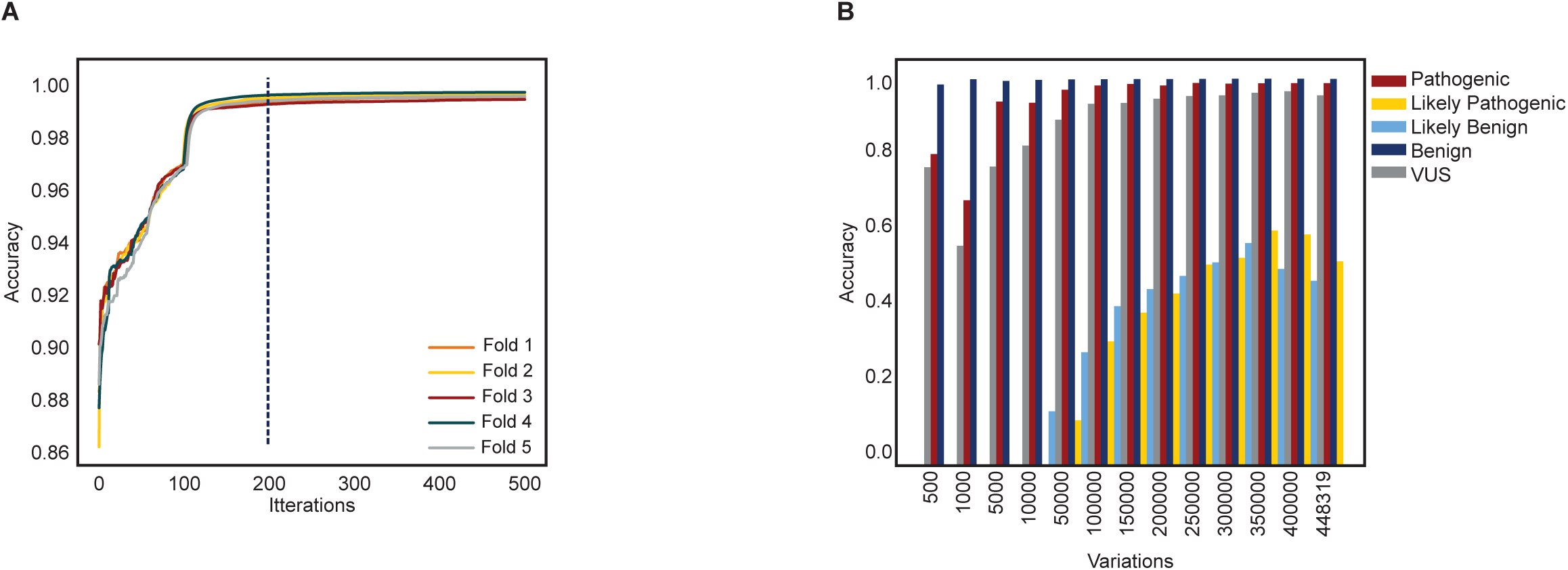
Azurify model parameterization. **A**: Testing accuracy (y-axis) shows training beyond 200 iterations (x-axis) does not markedly improve the accuracy of GBDT model (< 0.000001%). **B**: Variant sampling in the training data shows accurate classification (y-axis) can be rapidly achieved for pathogenic, benign and to a lesser extent VUS variants. However, accurate classification of less frequent labels in the training data (i.e. likely benign and likely pathogenic) require a large n (> 300000).

**Figure S3.**
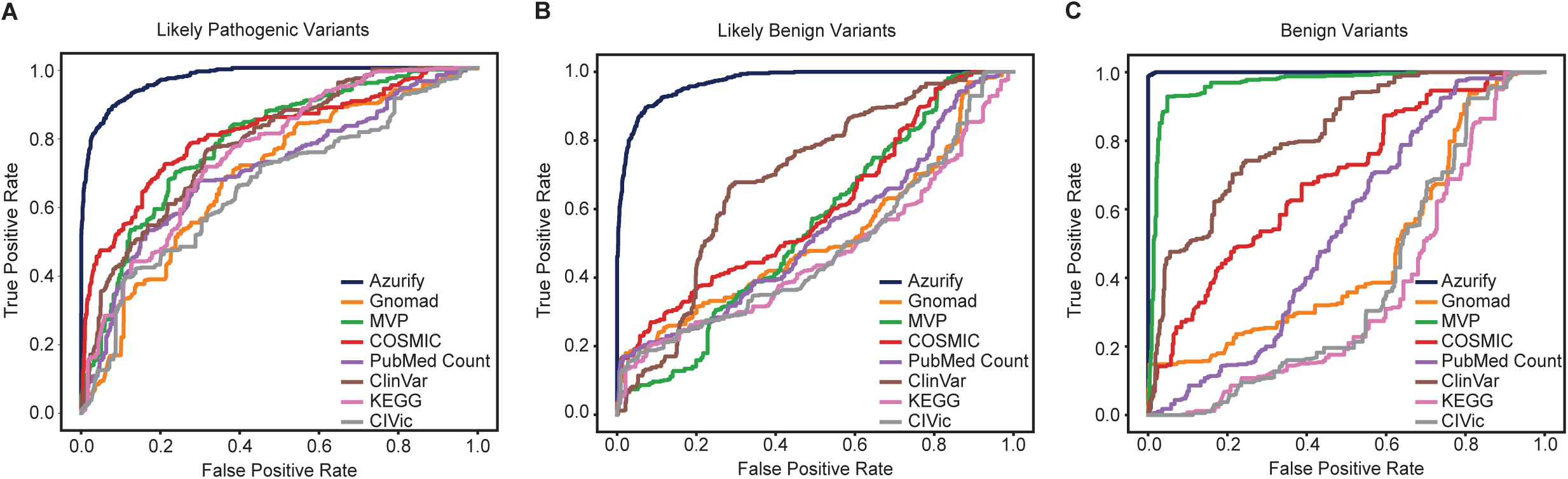
ROC Analysis of individual features by class. **A-C**: ROC curves of each feature’s ability to classify variants in the holdout data. Each ROC curve shows performance evaluation in predicting likely pathogenic (A), likely benign (B), and benign (C) variants when a given knowledgebase (i.e, gnomAD, MVP, etc) is added to genomic features (i.e, domain, coding effect, etc).

**Figure S4.**
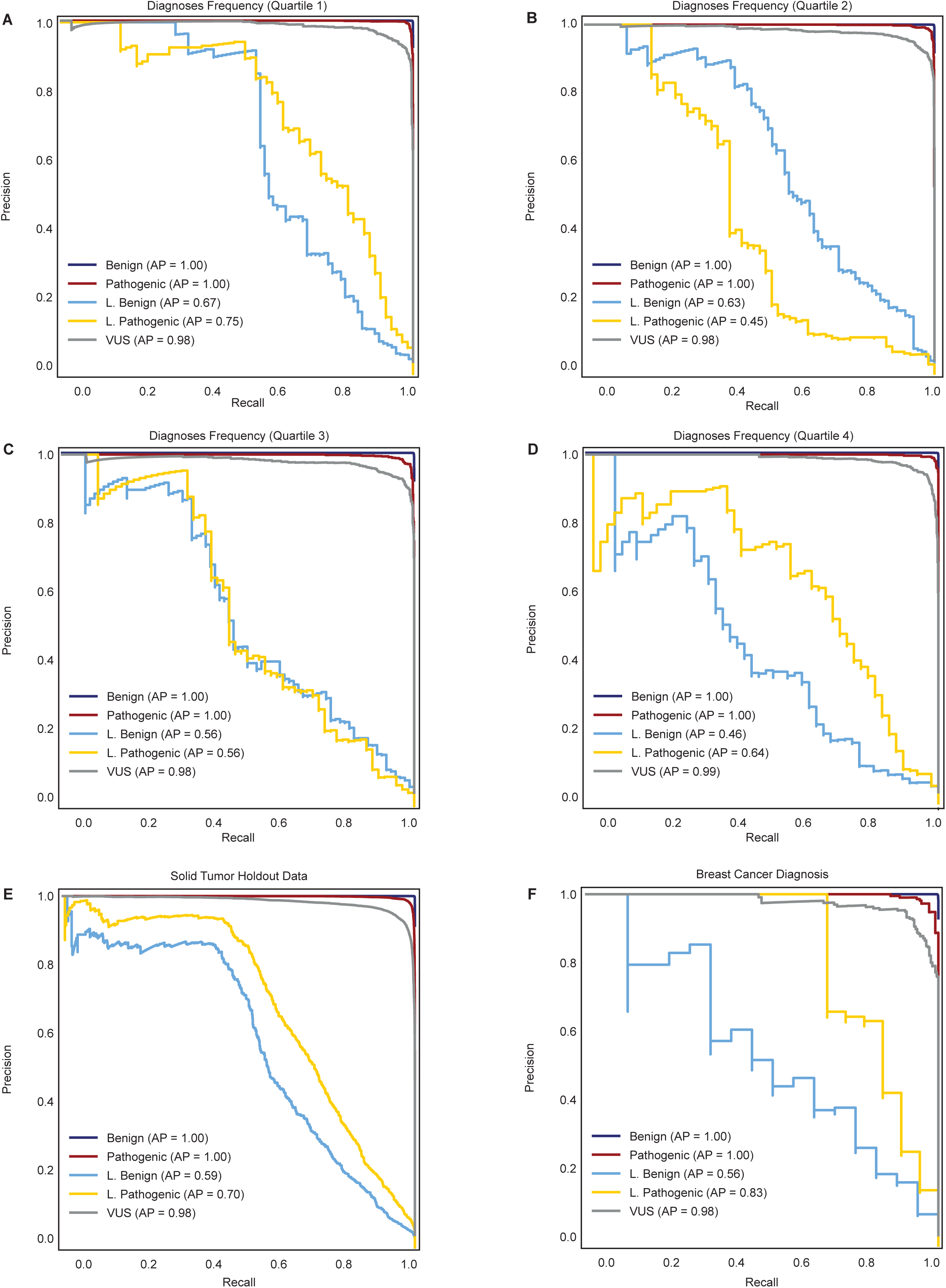
Precision-recall analysis based on solid tumor prevalence in the UPenn cohort. **A-D**: Precision-recall curves of solid tumor phenotypes, divided into quartiles based on presentation frequency in the holdout data. Each precision-recall curve shows performance in predicting all 5 Azurify classes in quartile 1 (A), quartile 2 (B), quartile 3 (C), and quartile 4 (D). **E-F**: Precision-recall curves for all the solid tumor (E) and only breast cancer (F) cases in the holdout data. This analysis shows no marked difference between Azurify average precision for classifying variants in breast cancer compared to other solid tumors in the UPenn solid tumor holdout cohort.

**Figure S5.**
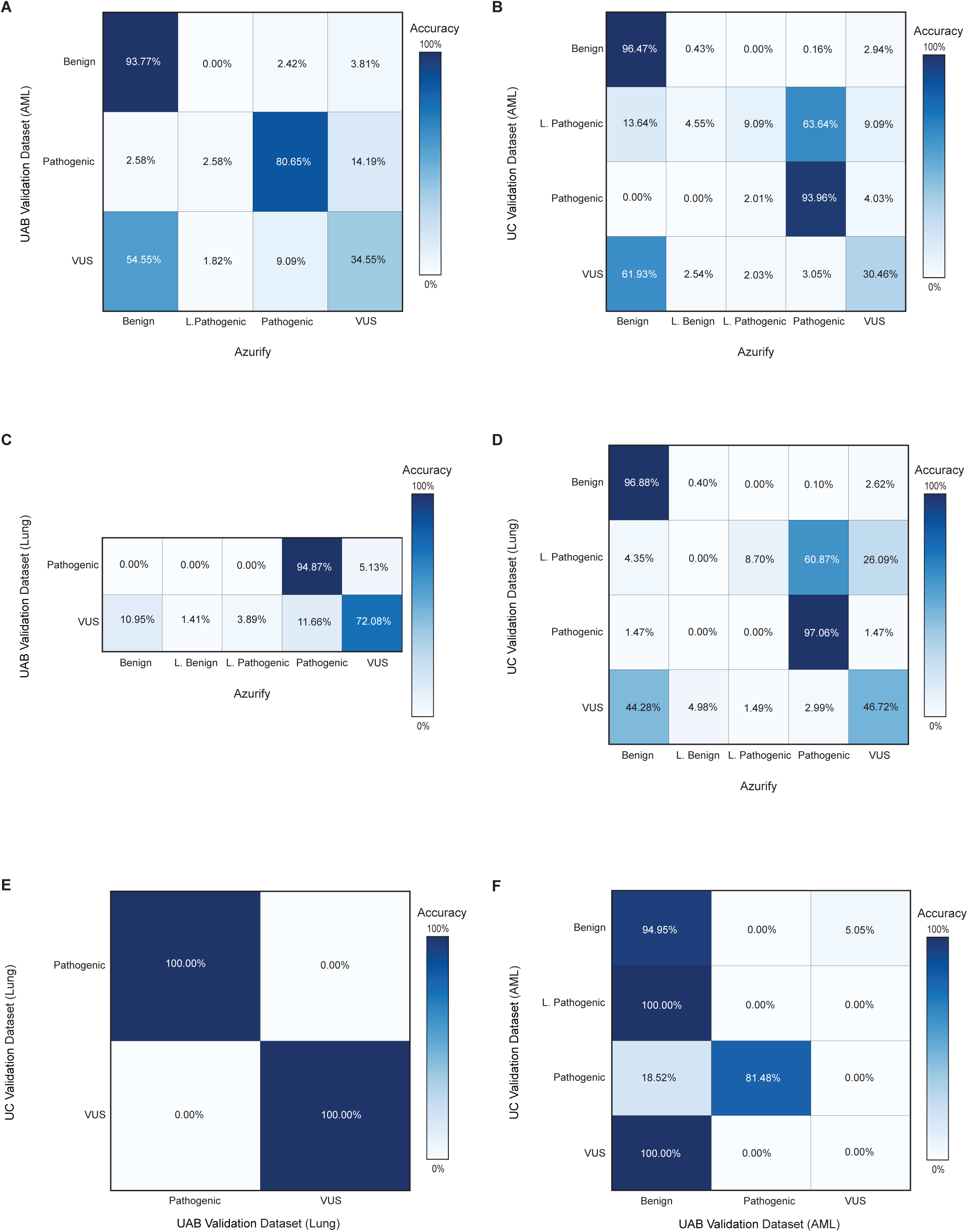
Azurify performance assessment with variants from lung cancer and AML cases UAB and UC CAP/CLIA-certified laboratories. **A**: Heatmap showing the Azurify classification accuracy for clinically reported somatic variants (n = 2,650) across 53 genes in 31 AML cases from UAB, where variants are graded into three classes: benign, pathogenic and VUS. **B**: Heatmap showing the Azurify classification accuracy for clinically reported somatic variants (n = 6,450) across 150 genes in 20 AML cases from UC, where variants are graded into four classes: pathogenic (Tier 1), likely pathogenic (Tier 2), VUS (Tier 3), and benign (Tier 4). **C**: Heatmap showing the Azurify classification accuracy for clinically reported somatic variants (n =361) across 211 genes in 30 lung cancer cases from UAB, where variants are graded into two classes: pathogenic and VUS. **D**: Heatmap showing the Azurify classification accuracy for clinically reported somatic variants (n = 5,254) across 158 genes in 20 lung cancer cases from UC, where variants are graded into four classes: pathogenic (Tier 1), likely pathogenic (Tier 2), VUS (Tier 3), and benign (Tier 4). **E:** Comparison of variant labels in UC and UAB lung cancer cases. Despite different variant reporting schema, classification of 31 mutually reported variants in 6 genes were 100% concordant. **F:** Comparison of variant labels in UC and UAB AML cases. Despite different variant reporting schema, classification of 954 mutually reported variants in 24 genes were concordant.

**Figure S6.**
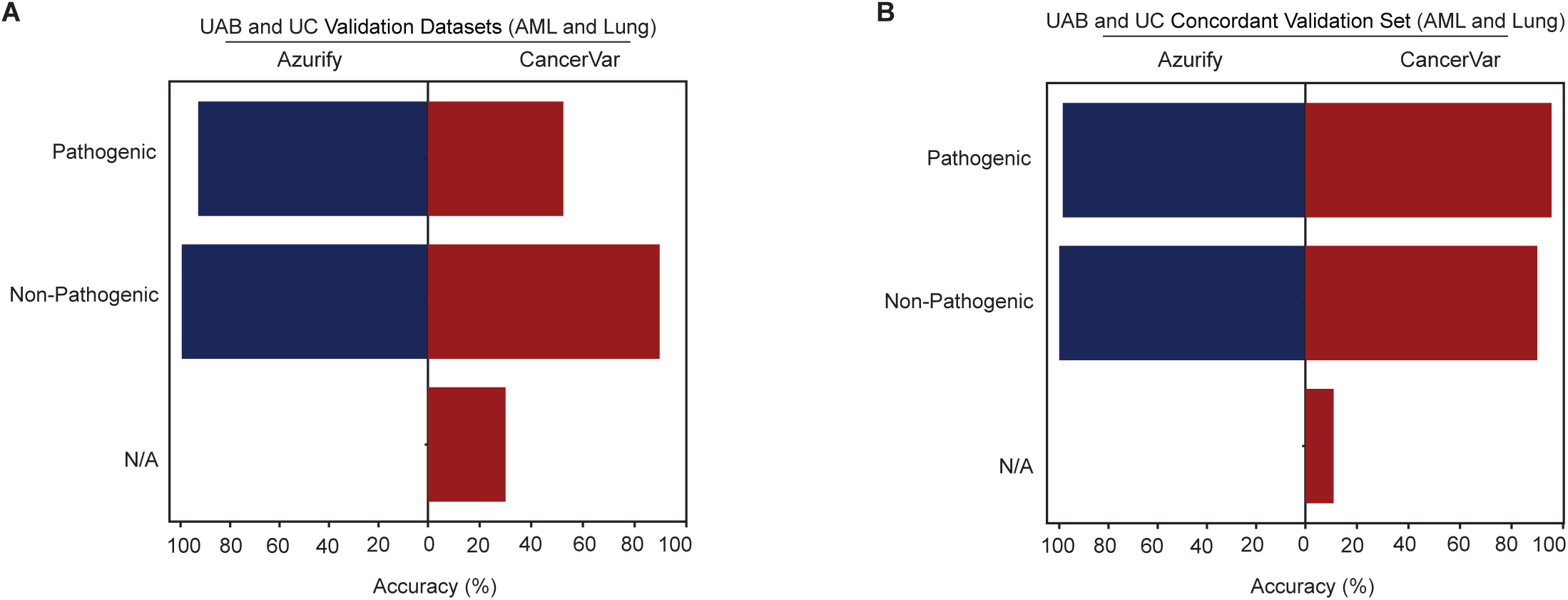
Azurify outperforms CancerVar in annotating data from UAB and UC CAP/CLIA-certified laboratories. **A**: Azurify (blue) and CancerVar (red) classification accuracy % for all the variants reported for AML and lung cancer cases by UAB and UC (n = 14,725). N/A: variants that an algorithm failed to classify. **B**: Azurify (blue) and CancerVar (red) classification accuracy % for variants concordantly reported for AML and lung cancer cases by UAB and UC (n = 985). N/A: variants that an algorithm failed to classify.

**Figure S7.**
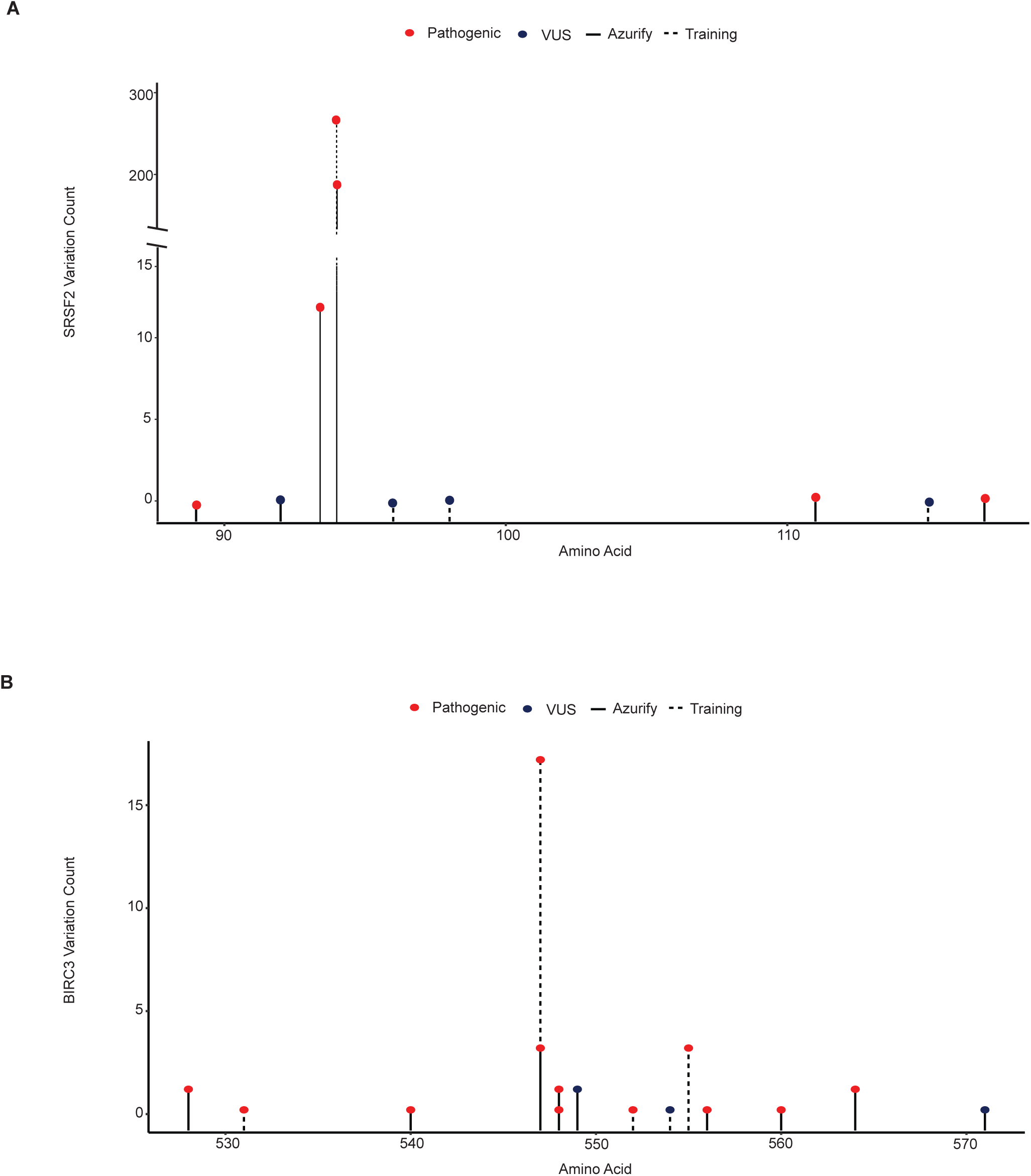
Azurify classifies variants in AML and CLL. **A**: Zoomed-in lollipop plot of SRSF2 mutations showing that Azurify is still able to accurately classify additional pathogenic and VUS variants around amino acid 95, a common variant in AML. Note that only a subset of pathogenic variants was present in the Azurify training data, as marked by dash lines. **B**: Zoomed-in lollipop plot of BIRC3 mutations showing that Azurify is still able to accurately classify pathogenic and VUS variants around amino acids 537-564, which are known to be pathologically relevant in CLL. Note that only a subset of pathogenic variants was present in the Azurify training data, as marked by dash lines.

## Supplemental Table Legends

**Table S1.** Cancer phenotypes used in the Azurify model training.

**Table S2.** Features and resources used in the Azurify model.

**Table S3.** Frequency of solid tumor phenotypes in the holdout data.

**Table S4.** Genes included in model training and UPenn independent datasets.

**Table S5.** UAB and UC meta-data and gene lists.

**Table S6.** Histology / disease count in the UPenn independent cohort.

